# The more you know: Investigating why adults get a bigger memory boost from semantic congruency than children

**DOI:** 10.1101/456624

**Authors:** Wei-Chun Wang, Simona Ghetti, Garvin Brod, Silvia A. Bunge

## Abstract

Humans possess the capacity to employ prior knowledge in the service of our ability to remember; thus, memory is oftentimes superior for information that is semantically congruent with our prior knowledge. This congruency benefit grows during development, but little is understood about neurodevelopmental differences that underlie this growth. Here, we sought to explore the brain mechanisms underlying these phenomena. To this end, we examined the neural substrates of semantically congruent vs. incongruent item-context associations in 116 children and 25 young adults who performed encoding and retrieval tasks during functional MRI data collection. Participants encoded item-context pairs by judging whether an item belonged in a scene. Episodic memory was then tested with a source memory task. Consistent with prior work, source memory accuracy improved with age, and was greater for congruent than incongruent pairs; further, this congruency benefit was greater in adults than children. Age-related differences were observed across univariate, functional connectivity, and multivariate analyses, particularly in lateral prefrontal cortex (PFC). In sum, our results revealed two general age differences. First, left ventrolateral/rostrolateral PFC exhibited age-related increases in univariate activity, as well as greater functional connectivity with temporal regions during the processing of congruency. Second, right rostrolateral PFC activation was associated with successfully encoded congruent associations in adults, but not children. Finally, multivariate analyses provided evidence for stronger veridical memory in adults than children in right ventrolateral PFC. These effects in right lateral PFC were significantly correlated with memory performance, implicating them in the process of remembering congruent associations. These results connect brain regions associated with top-down control in the congruency benefit and age-related improvements therein.

## Introduction

As we accumulate an understanding about the world around us, we use our semantic knowledge to help us learn and remember events in our daily lives. Indeed, it has long been recognized that humans possess the remarkable facility to employ prior knowledge in the service of our ability to remember (Bartlett, 1932), and that, subsequently, memory is typically superior for information that is semantically congruent with our prior knowledge (Craik & Tulving, 1975). Research shows that semantic knowledge supports both the elaboration and the organization of incoming information (Brod, Werkle-Bergner, & Shing, 2013; Ericsson & Kintsch, 1995), with consequent benefits for later episodic memory (Poppenk, Köhler, & Moscovitch, 2010; Reder et al., 2013; Wang, Brashier, Wing, Marsh, & Cabeza, 2018). This empirical evidence matches our experience: we can all recall experiences in which we more easily remember a news event that is related to a topic we are familiar with.

Along these lines, previous work also indicates that semantically congruous information (e.g., a deer in a park) is remembered better than semantically incongruous information (e.g., a cactus in a city) (Craik & Tulving, 1975; Cycowicz, Nessler, Horton, & Friedman, 2008; Staresina, Gray, & Davachi, 2008). This integration of to-be-remembered information with prior knowledge – or congruency benefit – allows for more elaborative encoding of associations and better subsequent memory. During typical development, this congruency benefit increases with age (Stangor & McMillan, 1992). A recent study (Brod, Lindenberger, & Shing, 2017) shows that this age-related increase likely reflects growth in semantic knowledge and/or an increase in the capacity to strategically associate information. By experimentally inducing prior knowledge to a comparable degree in children and adults and controlling for strategy use, 8–11 year-olds and adults demonstrated a comparable congruency benefit (Brod et al., 2017).

Here, we used functional MRI (fMRI) to better understand the mechanisms underlying the congruency benefit and why it improves over typical development. There has been nearly no research on encoding processes that may contribute to the congruency benefit in development, with the exception of one study examining differences between children (ages 8.5-11.5) and young adults during encoding (Maril et al., 2011; for differences during retrieval, see Brod et al., 2017). In the study by Maril and colleagues, children and adults encoded objects associated with typical and atypical colors (e.g., a red apple versus a purple dog). Young adults outperformed children in a subsequent item recognition test, but the congruency benefit was similar across age groups. An analysis of the neural substrates of this benefit revealed stronger activation for adults than children in prefrontal cortex (PFC) and parietal cortex regions associated with attention and control, as well as occipito-temporal regions associated with semantic memory; by contrast, children engaged occipital cortex to a greater extent than adults. These results suggest that the similar behavioral benefits may have resulted from different mechanisms, with adults relying more on control and semantic memory regions during the encoding of congruent associations and children on bottom-up, perceptual encoding processes. Maril et al. proposed that adults use their greater cognitive control capacity to orient their attention and selectively recruit relevant semantic knowledge to form congruent associations.

While this initial study provided an important insight into the neural underpinnings of the congruency benefit, it has some limitations. First, it was based on a small sample (15 children and 18 young adults). Second, the item recognition task may not have been as sensitive to age differences as a would be a task targeting retrieval of item-context associations (Ghetti & Bunge, 2012). Finally, the univariate analyses may not have identified age differences that could be observed with current multivariate and functional connectivity methods.

With the current fMRI dataset, we sought to investigate age differences in the neural correlates of the congruency benefit to better understand the development of semantic and episodic memory (see preregistration at http://aspredicted.org/blind.php?x=s5cq5y). Our study examined memory for item-context associations, a form of episodic memory for relational information. Given that the congruency benefit is dependent on the successful formation and retrieval of semantic associations, and that episodic memory improves during childhood (Ghetti & Angelini, 2008), this source memory task served as an ideal memory measure with which to examine semantic congruency effects.

We collected data on this task in a sample of 116 children (64 8-9-year-olds and 52 10-12-year-olds), along with 25 young adults between ages 18 and 25. We preregistered the intention to examine these younger and older child groups separately, because of prior research indicating that, during middle childhood, episodic memory performance improves rapidly (Ghetti & Angelini, 2008), along with associated changes in its neural correlates (DeMaster, Pathman, Lee, & Ghetti, 2013; Fandakova et al., 2016; Sastre III, Wendelken, Lee, Bunge, & Ghetti, 2016). Moreover, there is evidence that episodic memory following semantic compared to perceptual encoding is greater in older but not younger children (Ghetti & Angelini, 2008), thus raising the possibility that younger and older children may exhibit differences in the congruency benefit and its neural correlates. In addition to probing age differences, our large sample allowed us to assess individual differences. Investigating the neural correlates of the congruency benefit in development should help us to better understand why adults typically remember congruent information to a greater degree than children.

During encoding, one possibility is that adults, relative to children, draw on their more expansive semantic knowledge in order to form congruent associations, a process that recruits the semantic memory system (Binder, Desai, Graves, & Conant, 2009). We posited that such developmental differences would be reflected in: 1) greater univariate activation and/or 2) multivariate pattern similarity within brain regions implicated in semantic memory, and/or 2) greater functional connectivity among these regions.

Another possibility – not mutually exclusive with the first one – is that adults rely more strongly than children on fronto-parietal regions important for cognitive control to flexibly generate associations that are distinctive, but still semantically congruent, at encoding. We posited that such developmental differences would be reflected in 1) greater univariate/multivariate effects within control and attention regions, 2) greater functional connectivity between cognitive control and episodic memory regions, and/or 3) greater functional connectivity between cognitive control and semantic memory regions.

We tested these hypotheses using fMRI data from a memory task in which participants encoded items and contexts by judging whether each item “belonged” with one of three given scene contexts (e.g., whether a deer belonged in the city). At test, participants completed a source task for each item (i.e., which scene was this item paired with). We conducted three complementary analyses within relevant regions of interest (ROIs). These ROIs include regions implicated in semantic memory in a meta-analysis from Binder et al. (2009) (i.e., ventral parietal cortex, middle/inferior temporal, fusiform, and parahippocampal gyri, medial and ventrolateral PFC, and posterior cingulate gyrus). Many of these ROIs overlap with regions implicated in episodic memory. Additional ROIs also include regions that are more directly implicated in cognitive processes that support episodic memory: the hippocampus (Eichenbaum, Yonelinas, & Ranganath, 2007), dorsal parietal cortex (Cabeza, Ciaramelli, Olson, & Moscovitch, 2008; Uncapher & Wagner, 2009), dorsolateral PFC (Blumenfeld & Ranganath, 2007), and rostrolateral PFC (Simons & Spiers, 2003).

First, to identify brain correlates of the congruency benefit, we conducted voxel-wise analyses of univariate activation for Belong (i.e., congruent) compared to Don’t Belong (i.e., incongruent) trials at encoding within the above ROIs and tested individual differences related to behavioral measures episodic memory. Second, to explore interactions between brain regions during congruent processing, we examined functional connectivity between regions exhibiting significant univariate activation differences between Belong and Don’t Belong trials. Finally, we conducted pattern similarity searchlight analyses between encoding and retrieval trials within the abovementioned ROIs to assess whether congruency influenced the degree to which veridical memories are retrieved in the brain – and, if so, whether this effect differed as a function of age.

Together, this set of complementary analyses allowed us to investigate age-related differences in the congruency benefit and uncover the role of the processes driving the congruency benefit and the representational content related to congruent associations.

## Methods

### Participants

Behavioral and fMRI data were collected as part of a larger longitudinal study on memory development (Hippo Time; PIs Ghetti and Bunge) that has been described elsewhere (Fandakova et al., 2016; Fandakova et al., 2017; Wendelken et al., 2017). For the parent study, child participants were recruited primarily via flier distribution to elementary schools in the Sacramento City Unified School District in Sacramento, California, and the Washington Unified School District in West Sacramento, California. Young adults were college students recruited from the Department of Psychology’s subject pool at the University of California, Davis. Children received $70 for their participation; young adults received partial fulfillment of a course requirement at the University of California, Davis. Approval for study of human subjects was granted by the Institutional Review Board at the University of California, Davis.

For the present analyses, we examined data from all participants in the parent study for whom we had viable fMRI data at encoding for the item-context (or source) memory task, and who exhibited above-chance memory performance on the task (N = 141). Although the broader study was longitudinal, we used data from only the first timepoint in this initial characterization of developmental differences in the congruency benefit.

The final sample included 64 younger children (8.0–9.9 years, mean = 8.9 years, SD = 0.6 years, 30 females), 52 older children (10.0–12.0 years, mean = 10.7 years, SD = 0.5 years, 23 females), and 25 young adults (18.2–25.4 years old, mean = 19.3, SD = 1.4, 15 females). We selected the age groups based on previous studies which examined similar ages, both in overall range (DeMaster & Ghetti, 2013; DeMaster, Pathman, & Ghetti, 2013), and specific division into two child groups that have previously shown differences in brain activation related to episodic memory (DeMaster, Pathman, Lee, et al., 2013; Fandakova et al., 2016; Sastre III et al., 2016).

Data collected from 28 additional participants were excluded from the analyses. Five were excluded due to chance memory performance, and 23 were excluded as a result of excessive cutoff in the temporal or frontal lobes or excessive head motion during the scans (see **fMRI Data Analysis**). Specifically, scans with more than 25% bad volumes were excluded from analysis. Participants with at least two out of three usable retrieval scans based on this criterion were included. In addition, participants with only one usable retrieval scan were included if that scan had no more than 10% bad volumes. Within runs included in the final sample, 4.49% of possible volumes were excluded due to motion. Fewer volumes were excluded for adults (1.34%) compared to children (5.29%; p < .001).

### Materials and procedure

Data were collected at the UC Davis Imaging Research Center in Sacramento, California. All participants completed a brief training protocol using a mock scanner located at the imaging center. While in the scanner, participants completed a source memory task (see also, Fandakova et al., 2016; Fandakova et al., 2017; Sastre III et al., 2016). The task was subdivided into 3 alternating encoding and retrieval scans of 5 minutes and 6.5 minutes each, respectively.

During each encoding scan, participants viewed 48 item-context pairs and were asked to make a congruency judgment regarding the relation between the item and context (**Figure 1A**). On each trial, one of three contexts (city, park, or farm) was presented for 500 ms, followed by the appearance of a superimposed line-drawn item (an object or an animal; Cycowicz, Friedman, Rothstein, & Snodgrass, 1997) for 2000 ms. Participants were asked to assess whether or not that item “belonged” in that context and had 1000 ms to respond following the appearance of a “Does it belong?” prompt. Each trial was separated by a jittered fixation cross, ranging from 500 to 8000 ms in duration. Critically, items and contexts were selected to have no obvious associations, thus requiring participants to actively generate plausible relationships between them.

**Figure 1.**
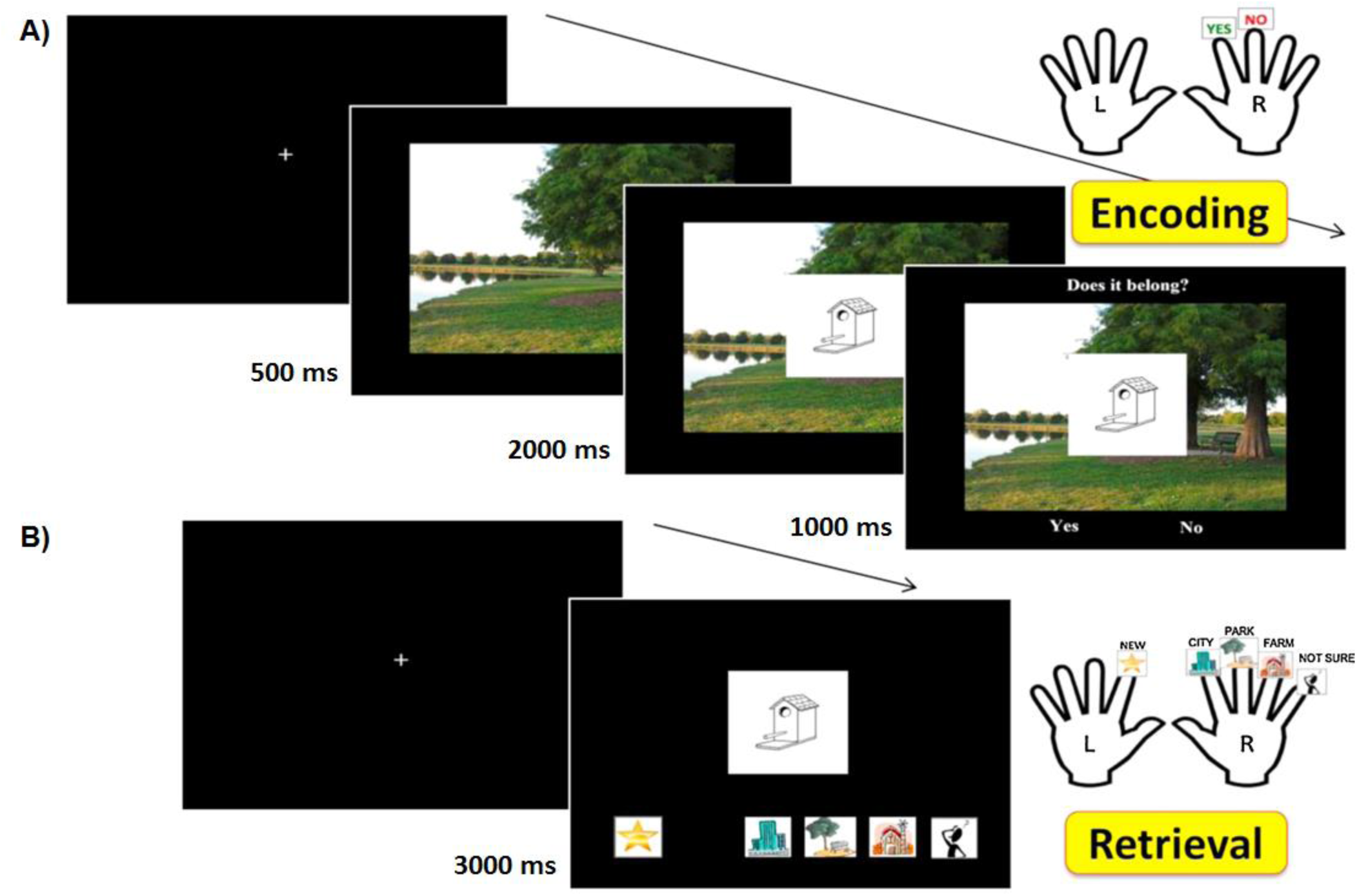
Participants completed (A) encoding and (B) retrieval phases of a source memory task in the scanner.

Participants were told to try to remember the item-context pairs for a subsequent memory test. The primary purpose of having them make the congruency judgment was to ensure that children and adults alike would pay attention and form associations, for our investigations of episodic memory development. The secondary purpose was to probe semantic memory development by examining behavioral and brain differences between item-context pairs that individual participants judged congruent (yes responses to the “Does it belong?” prompt) vs. incongruent.

For each retrieval scan (**Figure 1B**), participants were shown 64 line-drawn items, of which 16 were novel and 48 were items previously viewed in the preceding encoding scan. Participants were asked to either identify the item as novel, or identify the context with which it had been presented in the encoding scan (city, park, or farm). If participants remembered the item as previously seen, but could not remember the context with which it had been paired, they were instructed to select a “Don’t Know” button. This task was designed to be similar to source memory tasks used in previous studies (e.g., DeMaster & Ghetti, 2013; DeMaster, Pathman, & Ghetti, 2013; Güler & Thomas, 2013), but the use of scenes was thought to enrich the contextual information.

Participants completed the task inside the scanner using two 5-button LumiTouch button boxes, using the left-hand box for “new” responses and the right-hand box for all four “old” responses (i.e., city, park, farm, and “Don’t Know”). All participants were given a 5-min break outside the scanner after the first set of encoding and retrieval scans in order to reduce fatigue. The results of the retrieval phase are reported elsewhere (Sastre III et al., 2016; Selmeczy et al., in press)

### fMRI data acquisition

Each encoding and retrieval scan was completed inside the Siemens 3T MRI scanner using a 32-channel head coil. Functional MRI data were acquired with a gradient EPI sequence (repetition time (TR) = 2000 ms, echo time (TE) = 23 ms, no interstice gap, flip angle = 90°, field of view (FOV) = 204 mm, 148 volumes per scan for encoding and 196 volumes per scan for retrieval). Each volume consisted of 37 contiguous 3-mm axial slices. Voxel size was 3 mm × 3 mm × 3 mm. Foam padding was positioned between each participant's head and the coil to both ensure comfort and reduce head motion during the scan. All participants wore earplugs with a 29 db noise reduction rating to minimize scanner noise and facilitate communication with the experimenter.

### Behavioral analysis

Analysis of behavioral data included consideration of four separate measures: the congruency rate (i.e., the proportion of trials judged as Belong), item memory, source memory, and response times (RTs) at both encoding and retrieval (for RTs, see **Supplementary Results**). Memory and RT measures were calculated separately for Belong and Don’t Belong trials. Item memory was calculated as the hit rate, or the proportion of old items judged as old. Source memory was calculated as the number of hits with correctly identified source divided by the total number of hits (including “Don’t Know” responses).

### fMRI data analysis

Data were preprocessed and analyzed using SPM12 (http://www.fil.ion.ucl.ac.uk/spm/). Functional images were slice-time corrected, realigned, and coregistered to their respective anatomical images. The anatomical images were then segmented into separate grey and white matter images that were used to normalize the functional and anatomical images into MNI space. The normalization parameters were then applied to the functional images.

For univariate analyses, functional images were spatially smoothed with a 6-mm full-width half-maximum isotropic Gaussian kernel. The data were then high-pass filtered with a limit of 128 s and submitted to statistical analyses. Two separate encoding models were run in SPM12 using the general linear model (GLM). In the first model, Belong and Don’t Belong trials at encoding were modeled as conditions of interest. This allowed us to examine effects of congruency irrespective of subsequent memory. In the second model, Source Correct Belong and Source Correct Don’t Belong trials were modeled as conditions of interest (with Source Incorrect and Miss Belong and Don’t Belong trials also being modeled but not examined due to low trial counts).

For both models, whole-brain analyses were performed using a GLM that incorporated task effects (i.e., the trial types described above), session effects, and a general linear trend. The model also included six motion parameters as covariates of non-interest. In addition, to account for effects of subject motion, outlier volumes – those with motion in excess of 2 mm or signal change in excess of 2% – were obtained from the Artifact Detection Toolbox (http://www.nitrc.org/projects/artifact_detect) and spike regressors corresponding to those volumes were added as additional covariates of non-interest.

Task effects were modeled via epoch regressors aligned to the onset of each encoding trial, with the epoch duration equal to the encoding RT for that trial. Including RT in the model in this manner helped to minimize the extent to which increased RTs can drive increase BOLD signal (Grinband, Wager, Lindquist, Ferrera, & Hirsch, 2008). Parameter estimates associated with each trial type were combined to produce contrast images for target contrasts. Specifically, the difference between Belong and Don’t Belong trials (either for all trials or for Source Correct trials) was contrasted and analyzed in whole-brain one-way ANOVAs at the second level. Posthoc and exploratory region-of-interest (ROI) analyses were performed in MATLAB.

Task-based functional connectivity analyses were performed with the CONN toolbox (Whitfield-Gabrieli & Nieto-Castanon, 2012). The preprocessed data and SPM models (described above) were imported and run through CONN. The data were denoised by regressing out white matter and CSF signal and the covariates of non-interest (i.e., motion parameters and spike regressors), and applying linear detrending and a high pass filter (128 s). Spherical ROIs (9mm) were created from each significant cluster identified in the univariate analyses of all Belong vs. Don’t Belong trials, and then functional connectivity (i.e., Fisher-transformed Pearson correlation coefficients) was calculated between each pair of ROIs. These functional connectivity values were then submitted in a mixed-design ANOVA to assess effects of age group, congruency, and age group by congruency interactions.

To probe the representation of individual items, we performed an Encoding-Retrieval Similarity analysis on unsmoothed single trial beta values calculated using the least squares single (LSS) approach (Mumford, Turner, Ashby, & Poldrack, 2012). Encoding-retrieval similarity allowed us to examine reactivation of item-context associations across encoding and retrieval phases. A significant encoding-retrieval similarity effect is taken as evidence of the reactivation of specific items, as opposed to condition-level reactivation of the kind identified by multi-voxel pattern analysis (MVPA; Norman, Polyn, Detre, & Haxby, 2006); see **Supplementary Materials** for an MVPA analysis showing age-invariant reinstatement of encoding-related congruency processing at retrieval.

For item-level encoding-retrieval similarity, the encoding and retrieval activation patterns corresponding to the same trial were correlated, whereas for set-level encoding-retrieval similarity, encoding and retrieval activation patterns for each trial was correlated with the encoding activation patterns for all trials in that condition (Belong or Don’t Belong) and then averaged to create the whole-brain similarity volume for that retrieval trial. In the context of the present paradigm, if a deer were judged at encoding as belonging in a park, we would test for greater encoding-retrieval similarity with the deer at retrieval (i.e., item-level) than for all other items at retrieval (e.g., a ball) that were also judged as belonging with the scene they were paired with at encoding (i.e., set level).

Encoding-retrieval similarity was performed using an in-house searchlight script with a 3-voxel searchlight sphere (https://github.com/brg015/mfMRI_v2/) at both the item level and the set level. After encoding-retrieval similarity volumes were calculated for each retrieval trial, fixed-effect contrasts were generated separately for item-level and set-level encoding-retrieval similarity by averaging together all encoding-retrieval similarity volumes. These item-level and set-level volumes were then spatially smoothed (6-mm isotropic FWHM Gaussian filter) and submitted in mixed-design ANOVAs to assess effects of age group and encoding-retrieval similarity level (item vs. set) for Belong and Don’t Belong trials.

All univariate whole-brain and multivariate searchlight analyses were corrected for multiple comparisons with 3dClustSim (version 18.0.11) using an uncorrected threshold of p < .001 and a cluster extent of 40 voxels (for a discussion of cluster-level corrections, see Slotnick, 2017). Brain-behavior correlations were calculated with Pearson correlation coefficients. Both brain-behavior and functional connectivity correlations were FDR corrected to p < .05.

## Results

### Behavioral performance

#### Congruency proportion

To investigate age differences in the ability to meaningfully link items and contexts, we first examined the congruency proportion (**Table 1, top panel**), or the proportion of items judged as belonging to a given context, in a one-way ANOVA with our three age groups. This analysis revealed a significant effect of age group (F[2,141] = 7.19, p = .001, η_p_^2^ = .09). Post hoc t-tests confirmed that, compared to younger children, the congruency rate was greater for both young adults (p = .01) and older children (p = .001). The difference between older children and young adults was not significant (p = .81). Consistent with the interaction, the congruency proportion was significantly correlated with age among children (r = 0.22, p < .05). Thus, younger children were less likely to judge an item and context as being congruent.

**Table 1.** Congruency proportion, item memory, and source memory for each age group. SEM denoted in parentheses.

|  |  | Young Adults | Older Children | Younger Children |
| --- | --- | --- | --- | --- |
| Congruency Proportion |  | .45 (.03) | .46 (.02) | .38 (.02) |
| Item Memory<br>(Hit Rate) | Belong | .93 (.02) | .87 (.02) | .78 (.01) |
|  | Don't Belong | .88 (.03) | .81 (.02) | .73 (.02) |
|  | Difference | .05 (.01) | .05 (.01) | .05 (.02) |
| Source Memory<br>(Source Correct Rate) | Belong | .80 (.03) | .67 (.02) | .54 (.02) |
|  | Don't Belong | .60 (.03) | .55 (.02) | .41 (.02) |
|  | Difference | .21 (.03) | .11 (.02) | .13 (.02) |

#### Item memory

To examine age differences in the effect of congruency on memory, we first examined item memory (**Table 1, middle panel**) – or proportion of hits for items, regardless of source accuracy – as a function of age group and congruency. This mixed-design ANOVA revealed a significant main effect of age group (F[2,141] = 17.59, p < .001, ηp^2^ = .20). Post hoc tests indicated that item memory increased from younger to older children (p < .001) and from older children to adults (p < .05). In addition, there was a main effect of congruency (F[1,141] = 33.09, p < .001, ηp^2^ = .19), whereby item memory was greater for Belong than Don’t Belong trials. The interaction was not significant (F[2,141] = .07, p = .94, ηp^2^ = .001). Thus, congruency benefited item memory similarly in all age groups.

#### Source memory

We then tested whether congruency between associations at encoding would affect the capacity to subsequently remember associations. To do this, we examined source memory (**Table 1, bottom panel**), or the proportion of Source Correct responses as a function of age group and congruency. Similar to item memory, this mixed-design ANOVA revealed significant main effects of both congruency (F[1,141] = 132.94, p < .001, ηp^2^ = .49) and age group (F[2,141] = 25.57, p < .001, ηp_2_ = .27). These main effects, however, should be interpreted in the context of a significant two-way interaction (F(2, 141) = 3.63, p < .05, ηp^2^ = 0.05). Post hoc comparisons indicated that the congruency benefit was greater for young adults than either younger children (p < .05) or older children (p = .005), with no significant difference between younger and older children (p = .55). Together, these results suggested that the congruency benefit is greater in adults compared to children when remembering item-scene associations, but not items alone.

### fMRI analyses

#### Neural correlates of congruency processing

##### Univariate activation

To examine age-related differences in the processing of congruency independent of memory, we first examined brain activation at encoding, masked within regions previously associated with semantic and episodic memory (see **Methods**). A set of regions including bilateral occipital fusiform gyrus extending into parahippocampal cortex, middle/inferior temporal gyrus, left angular gyrus, and right superior parietal lobule exhibited greater activation for all Belong than Don’t Belong trials in a conjunction analysis of the three age groups (**Figure 2**, cool colors). In contrast, left ventrolateral/rostrolateral PFC exhibited an age group by congruency interaction (**Figure 2**, warm colors). A non-linear pattern was observed across the three age groups in this region, with greater activation differences for all Belong than Don’t Belong trials in young adults than children, and younger children than older children (see **Supplementary Figure 1**).

**Figure 2.**
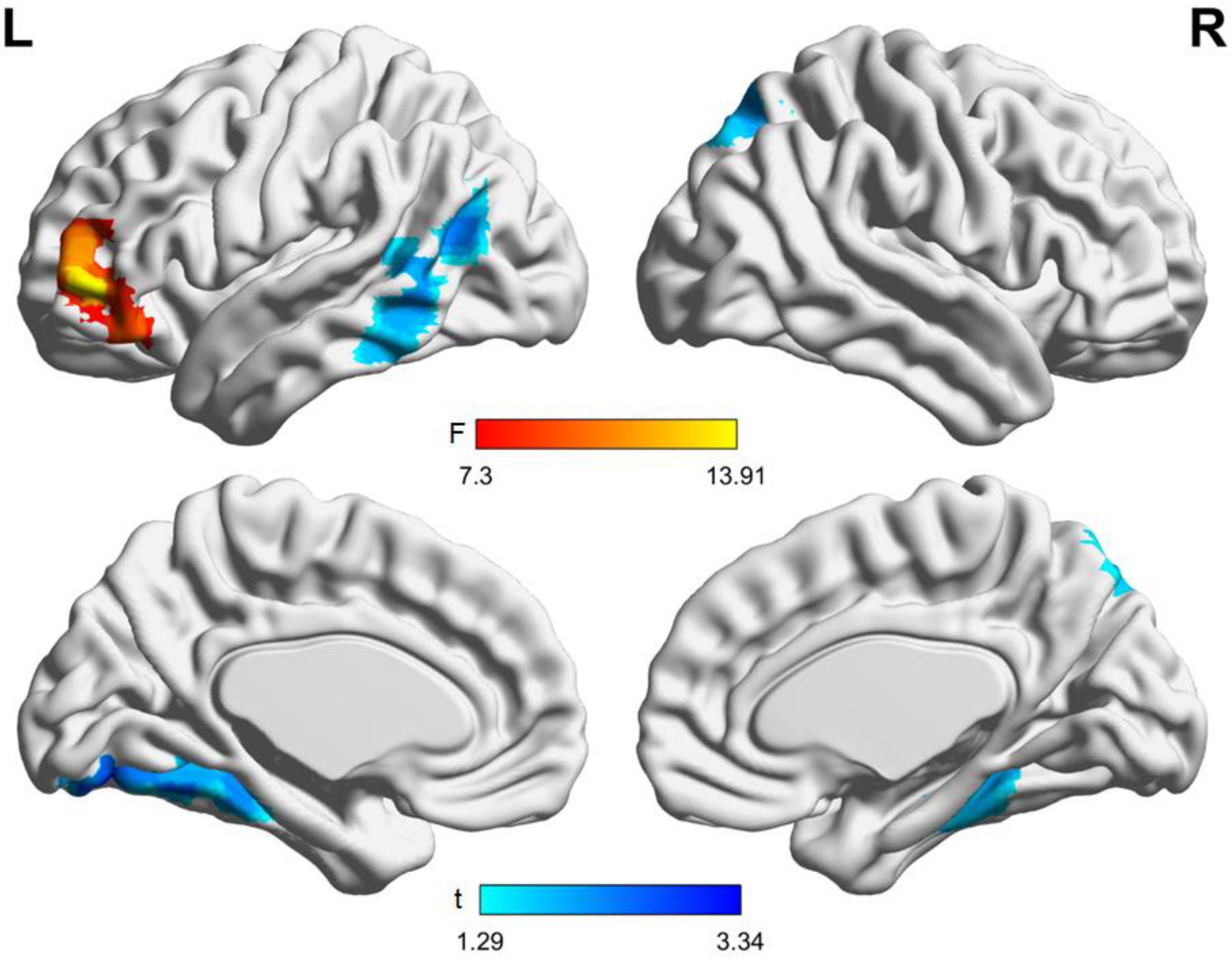
Regions exhibiting greater activation for Belong than Don’t Belong trials in a conjunction of the three age groups (cool colors) and exhibiting an age group by congruency interaction (warm colors).

##### Functional connectivity

Next, to further illuminate developmental differences in the processing of congruency, we tested for age-related differences in patterns of functional connectivity among the clusters that emerged from the univariate analysis of all Belong and Don’t Belong trials. To this end, we created six spherical ROIs, centered on the peak voxel of activation within the clusters reported above (also see **Supplementary Table 2**). We then tested for main effects of age group and condition, as well as age group by condition interactions. As illustrated in **Figure 3**, we found main effects of age group and interactions between age group and congruency.

**Table 2.** Significant functional connectivity effects of age group (after FDR correction) between regions exhibiting univariate effects. *Significant post-hoc t-test (p < .05). YA: Young adults; OC: Older children; YC: Younger children; vlPFC: ventrolateral/rostrolateral PFC; FG: Fusiform Gyrus; MTG: Middle Temporal Gyrus; SPL: Superior Parietal Lobule

| Contrast | F | Regions | Connectivity Differences (*t*) |  |  |
| --- | --- | --- | --- | --- | --- |
|  |  |  | YA vs. OC | YA vs. YC | OC vs. YC |
| Effect of Age Group | 7.13 | SPL ↔ L FG | 3.55* | 3.18* | -1.05 |
|  | 5.12 | SPL ↔ R FG | 2.91* | 2.68* | -0.47 |
|  | 3.98 | vlPFC ↔ R FG | 0.84 | 2.63* | 2.10* |
|  | 12.12 | MTG ↔ L FG | -2.17* | -4.40* | -3.21* |

**Figure 3.**
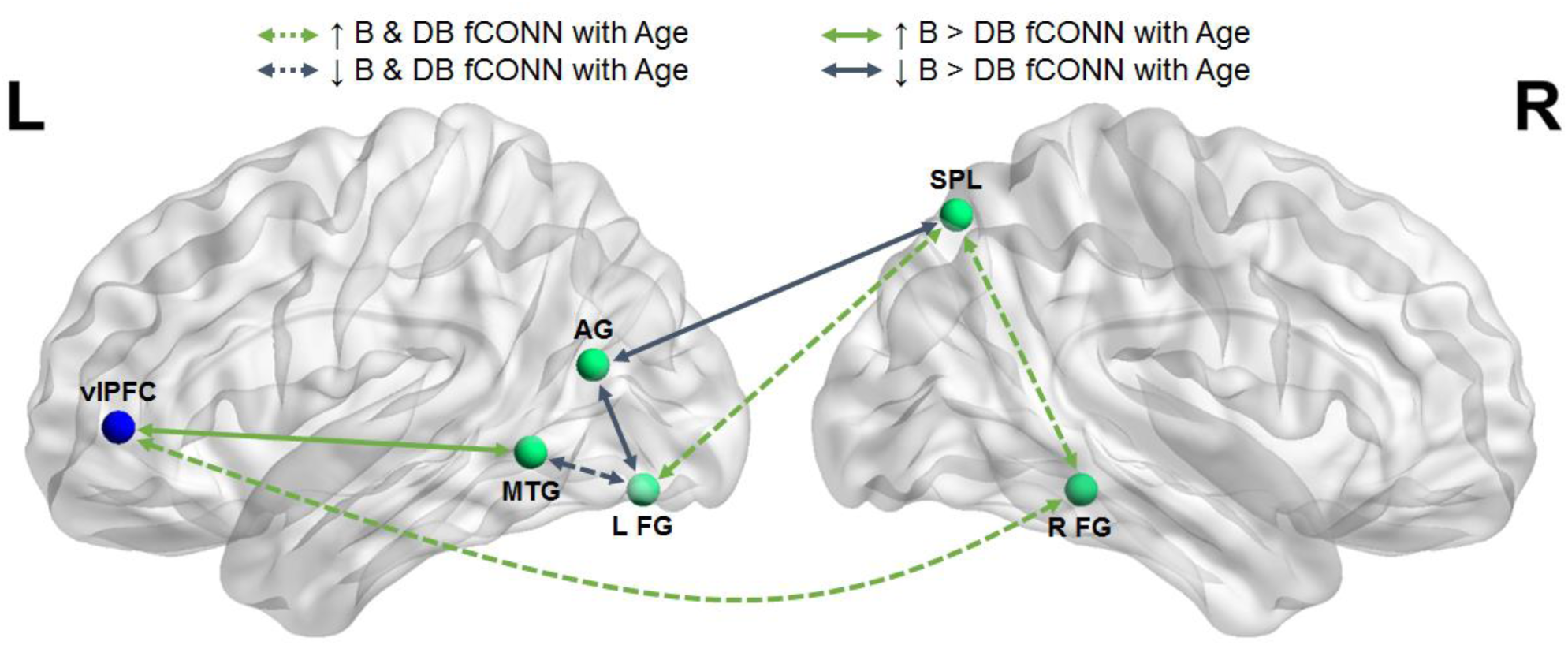
Functional connectivity between regions exhibiting univariate effects and interactions with group for Belong vs. Don’t Belong trials. vlPFC: ventrolateral/rostrolateral PFC; MTL: middle temporal gyrus; AG: angular gyrus; FG: fusiform gyrus; SPL: superior parietal lobule.

Regarding main effects of age group, we observed three distinct patterns (**Table 2**). First, connectivity between right superior parietal lobule and both left and right fusiform gyrus was significantly greater in young adults than both younger and older children (ps < .01), but did not differ between the child groups (ps > .30). Second, connectivity between left ventrolateral/rostrolateral PFC and right fusiform gyrus was significantly greater in both young adults and older children than younger children (ps < .05), but did not differ between young adults and older children (p = .36). The final main effect was found between left middle temporal and fusiform gyrus, which exhibited decreasing connectivity with age; young adults exhibited less connectivity than older children (p < .05), and older children less than younger children (p < .01).

Regarding age group by congruency interactions (**Table 3**), left ventrolateral/rostrolateral PFC and middle temporal gyrus (**Figure 4A**) exhibited increasing connectivity differences for Belong than Don’t Belong trials in young adults compared to both older and younger children (ps < .05), and did not differ between older and younger children (p = .14). On the other hand, left angular gyrus and both left fusiform gyrus (**Figure 4B**) and right superior parietal lobule (**Figure 4C**) exhibited a greater Belong than Don’t Belong connectivity difference in younger children compared to young adults (ps < .01), but did not differ between older children and either younger children or young adults (ps > .05).

**Table 3.**
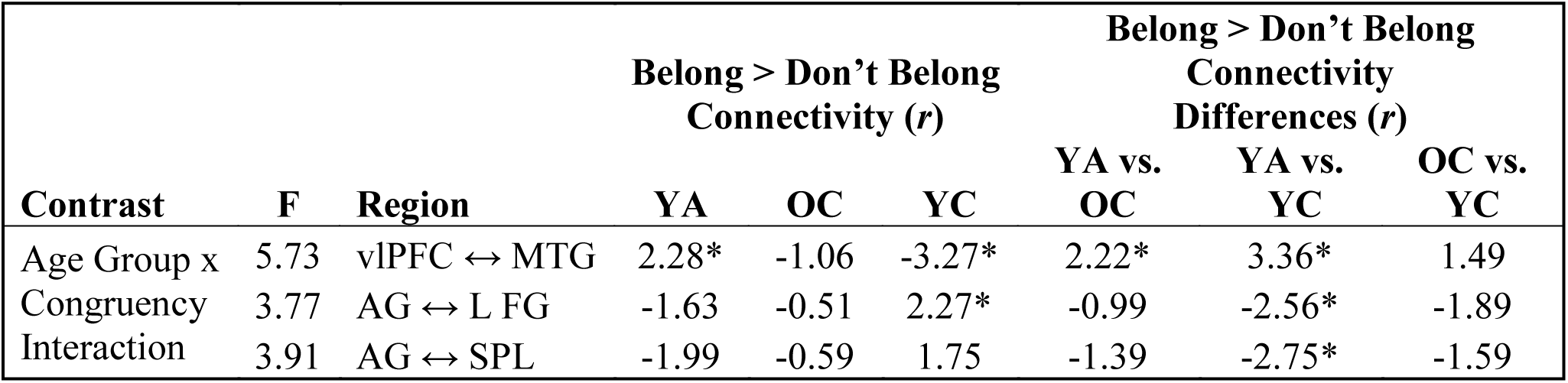
Significant functional connectivity age group by congruency effects (after FDR correction) between regions exhibiting univariate effects. *Significant post-hoc t-test (p < .05). YA: Young adults; OC: Older children; YC: Younger children; AG: Angular Gyrus; vl/rlPFC: ventrolateral/rostrolateral PFC; FG: Fusiform Gyrus; MTG: Middle Temporal Gyrus; SPL: Superior Parietal Lobule

**Figure 4.**
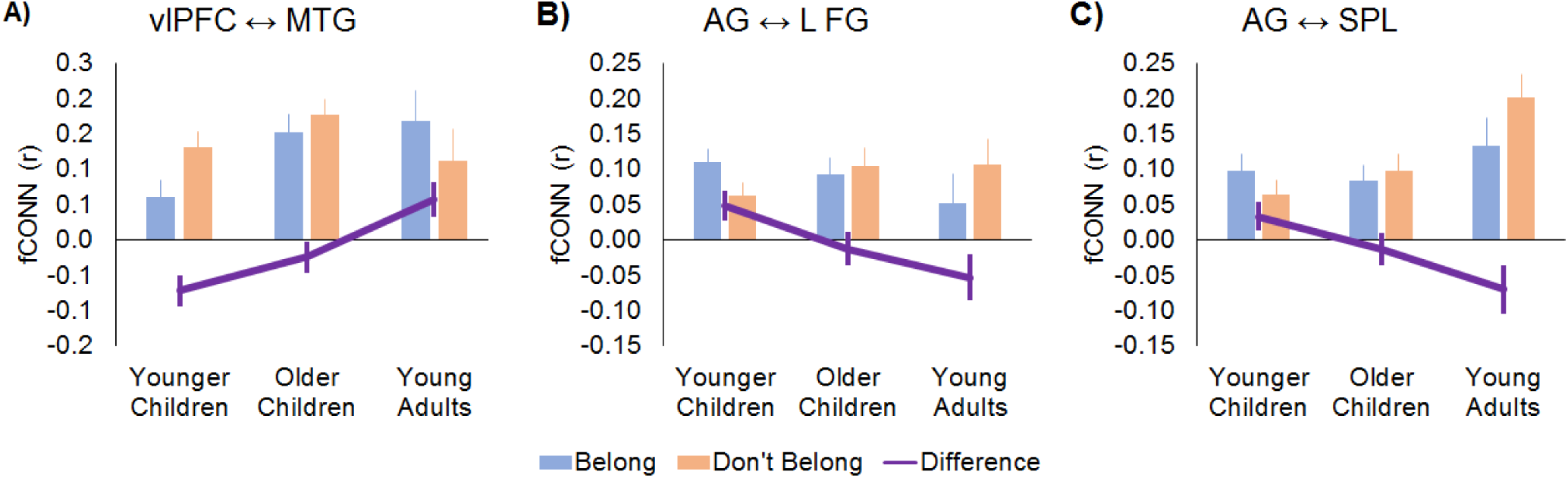
(A) Increasing Belong > Don’t Belong connectivity difference with increasing age group between ventrolateral/rostrolateral PFC (vlPFC) and middle temporal gyrus (MTG). (B) Decreasing Belong > Don’t Belong connectivity with increasing age group between angular gyrus (AG) and left fusiform gyrus (L FG). (C) Decreasing Belong > Don’t Belong connectivity with increasing age group between AG and superior parietal lobule (SPL).

Thus, across both Belong and Don’t Belong trials, we observed age-related decreases in connectivity between left fusiform and middle temporal gyrus, and age-related increases in connectivity between fusiform and frontoparietal regions. In addition, we observed age differences related to congruency: specifically, greater posterior brain connectivity for Belong than Don’t Belong trials in younger children, and greater anterior brain connectivity for Belong than Don’t Belong trials in young adults. Together, these results provide evidence for large-scale shifts in connectivity across age groups.

#### Univariate encoding activation associated with the congruency benefit

In the previous section, we examined the neural correlates of congruency processing to understand age differences in the ability to meaningfully integrate prior knowledge. Next, we sought to test how activation differences between Belong and Don’t Belong trials at encoding related to the congruency benefit. To this end, we conducted a whole-brain one-way ANOVA of univariate activation differences between source correct Belong and Don’t Belong trials, masked within regions previously associated with semantic and episodic memory. In a conjunction of the three age groups, no common clusters were found. By contrast, an F-contrast testing for age differences in activation revealed a significant cluster in right rostrolateral PFC (Z = 4.39, x = 48, y = 53, z = 2, k = 40; **Figure 5**). An ROI analysis revealed that young adults had greater Belong than Don’t Belong Source Correct activation in this region than both older and younger children (ps < .001), but there was no difference between younger and older children (p = .75). The difference in activation between Belong and Don’t Belong Source Correct trials was significant in young adults (p = .001), but not children (ps > .11). This lack of a difference in children was also reflected in a nonsignificant correlation between age and the magnitude of the activation difference in children (r = .12, p = .19). Finally, an exploratory analysis revealed a modest but significant correlation between activation and source memory differences between Belong and Don’t Belong trials across all participants (r = .20, p < .05). Taken together, these results suggest that right rostrolateral PFC contributes to the congruency benefit for source memory across participants, but that the effect is stronger for adults. Thus, right rostrolateral PFC activation reflects the congruency benefit for source memory, as well as the age-related increase in this congruency benefit.

**Figure 5.**
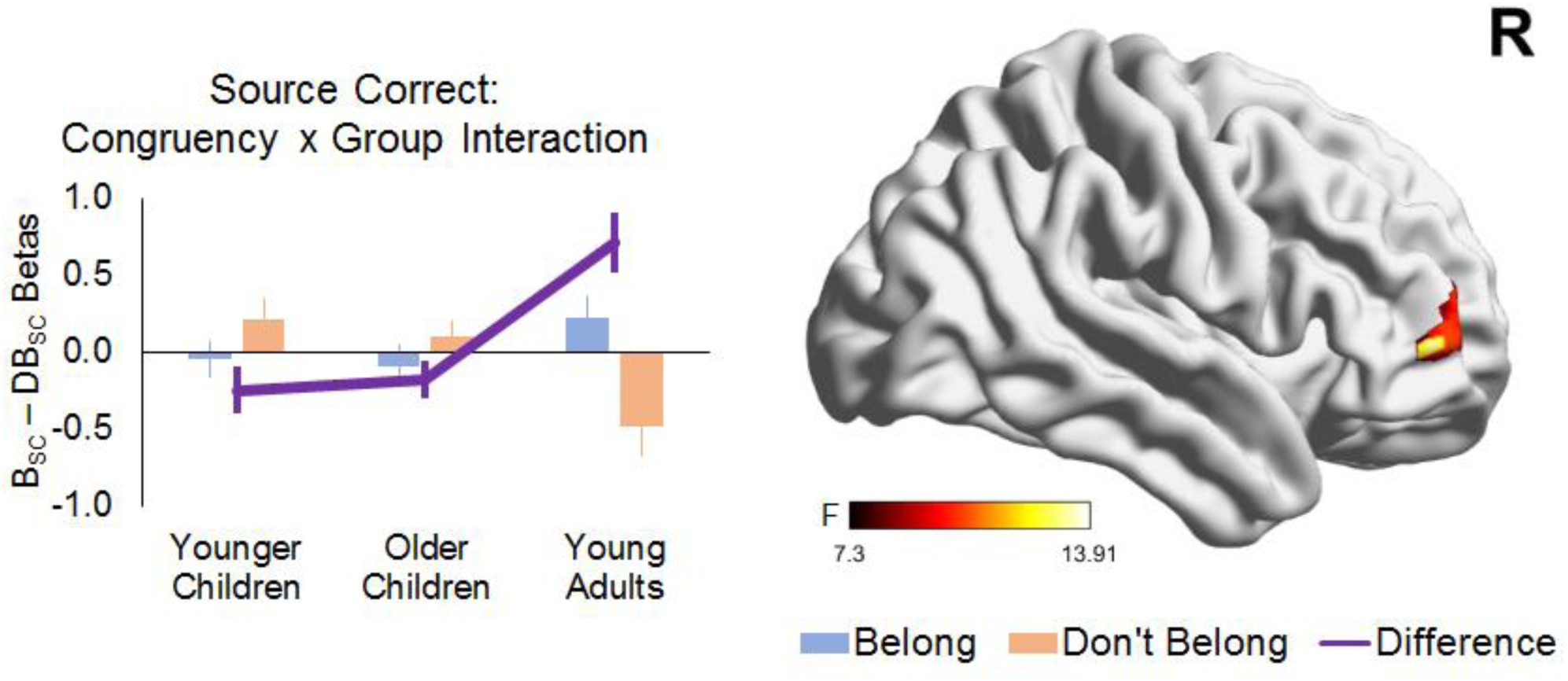
Right rostrolateral PFC exhibited greater Belong than Don’t Belong activity for Source Correct trials in young adults compared to both older and younger children.

#### Encoding-retrieval similarity

Lastly, we sought to investigate whether congruency influenced the degree to which veridical memories are retrieved in the brain – and, if so, whether this effect differed as a function of age. To this end, we examined voxel-wise activation patterns across encoding and retrieval scans, and tested whether the degree of pattern similarity differed as a function of congruency, memory, and/or age. To address these questions, we conducted an exploratory encoding-retrieval similarity analysis (Ritchey, Wing, LaBar, & Cabeza, 2012; Wing, Ritchey, & Cabeza, 2015). Encoding-retrieval similarity, which has never been assessed in children, allowed us to assess item-level similarity (i.e., activation patterns that are common to encoding and retrieval for specific items) relative to set-level similarity (i.e., activation patterns that are common to encoding and retrieval when the items themselves differ).

Our first analysis tested for an encoding-retrieval similarity effect (i.e., item-greater than set-level encoding-retrieval similarity), irrespective of age group or congruency. We found significant encoding-retrieval similarity in bilateral occipitotemporal cortices in a conjunction of the three age groups for both Belong and Don’t Belong trials (**Figure 6**), likely reflecting the reactivation of item-specific information. This is the first evidence of comparable item-level reactivation between adults and children using encoding-retrieval similarity. No suprathreshold clusters were observed when directly contrasting Belong vs. Don’t Belong trials (i.e., effect of congruency), and an F-contrast of age group also revealed no significant effects.

**Figure 6.**
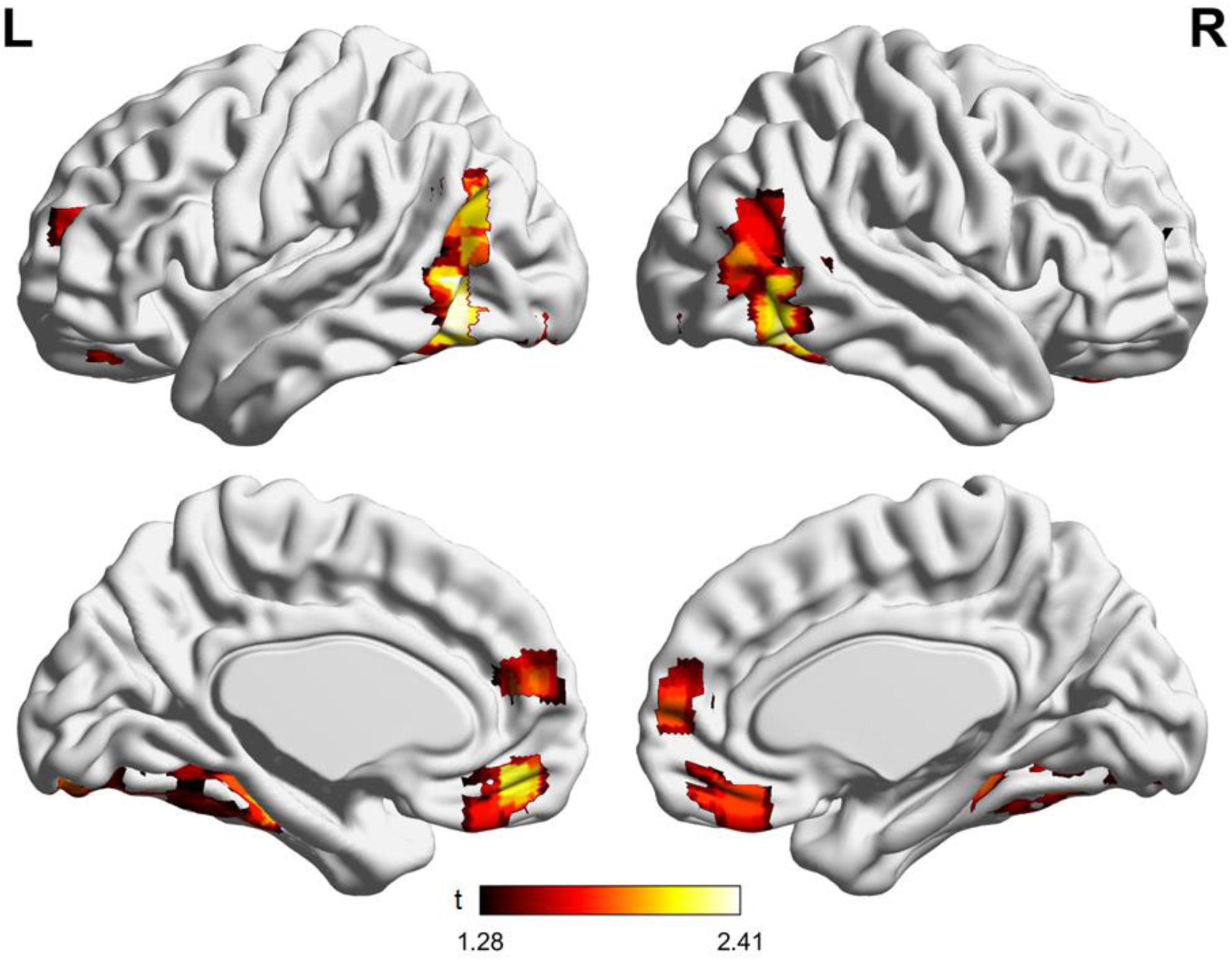
Regions exhibiting significant encoding-retrieval similarity (ERS) in both Belong and Don’t Belong trials in a conjunction of the three age groups.

Next, we searched for evidence of age group by encoding-retrieval similarity level interactions for Belong or Don’t Belong trials. For Belong trials, we observed a significant interaction, as measured by an F-contrast, in right ventrolateral PFC (Z = 4.31, x = 54, y = 35, z = -7, k = 71; **Figure 7A**). A follow-up ROI analysis of this cluster revealed that encoding-retrieval similarity was significantly above chance for young adults (p < .001), but not children (ps > .65). Moreover, encoding-retrieval similarity for young adults was greater than both younger and older children (ps < .001). Consistent with this finding, the magnitude of encoding-retrieval similarity did not significantly correlate with age among children (r = .09, p = .35). On the other hand, encoding-retrieval similarity in this ventrolateral PFC cluster correlated significantly with source memory for Belong Trials across all participants (r = 0.27, p < .01; **Figure 7B**). Additionally, the pattern of encoding-retrieval similarity in this region was similar when restricted to source correct trials only. In contrast, no regions exhibited a significant group by encoding-retrieval similarity interaction for Don’t Belong trials. Thus, while posterior regions exhibited age-invariant encoding-retrieval similarity effects, ventrolateral PFC exhibited greater encoding-retrieval similarity for Belong trials in young adults, mirroring the behavioral congruency benefit.

**Figure 7.**
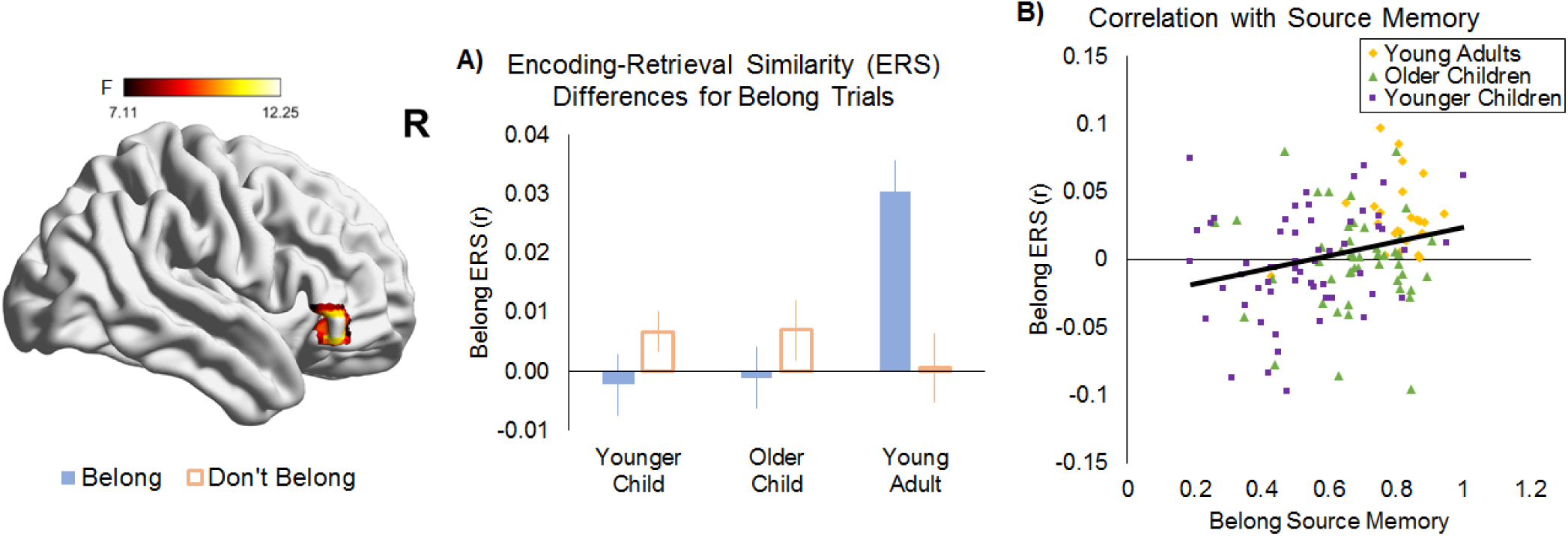
Regions exhibiting significant age group differences in encoding-retrieval similarity (ERS) for Belong trials. (A) Right ventrolateral PFC exhibited greater ERS for Belong trials in young adults compared to younger and older children. ERS for Don’t Belong trials is included for illustrative purposes. (B) For Belong trials, the magnitude of ERS and Source Memory was significantly correlated across all participants. For illustrative purposes, each age group is plotted separately.

Moreover, the magnitude of encoding-retrieval similarity correlated with source memory performance.

## Discussion

In the current study, we adopted three complementary analytic approaches to examine the neural substrates of the congruency benefit, and its growth during childhood. Behaviorally, we found that 10-12-year-olds and young adults reported a greater proportion of congruent responses than 8-9-year-olds, and that young adults exhibited a larger congruency benefit (i.e., better source memory for congruent than incongruent items) than both groups of children. This result, obtained in the context of an episodic memory paradigm, contrasts with the age-invariant congruency effects reported previously for item recognition (Maril et al., 2011). This finding suggests a graded process in typical development, wherein younger children are less capable of forming meaningful associations between unrelated pairs, and both younger and older children are less able to use successfully formed congruent associations to support their episodic memory.

We found age-invariant effects in temporal and parietal regions associated with semantic memory, and age differences in PFC regions associated with controlled semantic retrieval and elaboration, with greater congruency-related effects in young adults. First, young adults exhibited greater congruent than incongruent activation for all trials in left ventrolateral/rostrolateral PFC, and source correct trials in right rostrolateral PFC. Second, congruency-related functional connectivity shifted with age, increasing between left ventrolateral/rostrolateral PFC and middle temporal gyrus but decreasing between left angular gyrus and both left fusiform gyrus and right superior parietal lobule. Finally, young adults exhibited greater congruency-related encoding-retrieval similarity in right ventrolateral PFC; moreover, the magnitude of this reactivation correlated with source memory, implicating this region in congruent memory representations. These results suggest that developmental differences in lateral PFC may account for age-related improvements in our ability to both generate and remember congruent information.

### Age-related differences in congruency processing

Whereas temporal and parietal cortices associated with semantic memory exhibited age-invariant univariate effects, age-related univariate differences were revealed in lateral PFC. Across all trials, left ventrolateral/rostrolateral PFC exhibited a non-linear trend with age group: young adults showed a greater congruency-related effect (i.e., Belong > Don’t Belong) than both older and younger children, while younger children exhibited a greater congruency-related effect than older children. A greater congruency effect in lateral PFC in young adults is consistent with the slow maturation of this brain region (Huttenlocher & Dabholkar, 1997; Petanjek et al., 2011; Sowell et al., 2003). On the other hand, a greater congruency effect in young compared to older children was unexpected; one possibility is that the non-linear patterns of univariate activation across age groups are the result of the reorganization of large-scale brain networks during the transition from childhood to adulthood (Grayson & Fair, 2017).

Consistent with this possibility, we observed age differences in congruency-related functional connectivity. The fact that the pattern of stronger connectivity between congruent and incongruent trials decreased with age between parietal control regions and occipital fusiform gyrus, but increased between left ventrolateral/rostrolateral PFC and middle temporal gyrus, further suggests age-related increases in involvement in PFC regions, and may in part explain the presence of non-linear age-related differences in activation. This finding is consistent with previous work suggesting that during memory encoding, children rely more on perceptual features, whereas adults rely more on conceptual features (Brod et al., 2013; Maril et al., 2011; Ofen & Shing, 2013). Together, differences in congruency-related univariate activation and functional connectivity in left ventrolateral/rostrolateral PFC suggests that there is a shift toward frontal regions implicated in controlled semantic elaboration (Badre & Wagner, 2007; Bunge, Wendelken, Badre, & Wagner, 2004; Souza, Donohue, & Bunge, 2009; Wagner, Bunge, & Badre, 2004; Wagner, Paré-Blagoev, Clark, & Poldrack, 2001) from childhood to adulthood. This shift may be reflected in a reduction congruency-related univariate activation in older children relative to younger children and young adults, as well as a monotonic increase in congruency-related functional connectivity with age.

### Age-related differences in memory for congruent pairs

When we restricted our analyses to source-correct trials, we found an age-related difference in right rostrolateral PFC. Young adults, but not children, showed greater univariate activation for congruent than incongruent source-correct responses. While not typically associated with controlled semantic retrieval or elaboration, studies have found greater right rostrolateral PFC activation when judging the relatedness of concrete compared to abstract words (Sabsevitz, Medler, Seidenberg, & Binder, 2005) and when processing related compared to unrelated objects during memory encoding (Hawco, Armony, & Lepage, 2013). Speaking to its importance for successful memory encoding, a previous study found a greater subsequent memory effect in right rostrolateral PFC in associations for which prior knowledge was of high compared to low relevance (Brod, Lindenberger, Wagner, & Shing, 2016). Our results, along with these prior studies, suggest a role for right rostrolateral PFC in the elaboration of associations during successful memory encoding. Moreover, in line with work suggesting that memory-related effects in PFC increase with age (Brod et al., 2017; Shing, Brehmer, Heekeren, Bäckman, & Lindenberger, 2016; Tang, Shafer, & Ofen, 2017), this result suggests that maturation of rostrolateral PFC leads to an increase in its role in the successful encoding of congruent associations. Lastly, across all participants, activation differences between Belong and Don’t Belong source-correct trials significantly correlated with source memory, further suggesting that this region contributes to the successful encoding of congruent associations.

In addition to examining univariate activation differences, we also conducted an exploratory encoding-retrieval similarity analysis, to examine trial-specific reactivation across encoding and retrieval phases. Consistent with prior work (Wing et al., 2015), we observed reactivation for both congruent and incongruent trials in lateral and ventral temporal regions. While we did not observe differences related to congruency, right ventrolateral PFC exhibited greater reactivation for Belong trials in young adults compared to both younger and older children. Moreover, magnitude of this reactivation for Belong trials correlated with source memory accuracy, suggesting that right ventrolateral PFC plays a selective role in representing congruent associations that are successfully encoded. While previous studies suggest that PFC regions are critical for control processes important for memory encoding, our results are consistent with recent work suggesting that these regions may also contain representation content (e.g., Long & Kuhl, 2018), and to our knowledge this is the first study to examine encoding-retrieval representational differences in development.

### Limitations

One important limitation of the current study is that many participants contributed too few source incorrect responses to assess differences between source correct and incorrect trials, a standard way of assessing memory differences in fMRI. However, given our interest in the congruency benefit, as well as the cognitive process of judging congruency, our comparison of source correct trials is still informative. Moreover, regions exhibiting age differences related to memory– either univariate or representational differences – exhibited significant correlations with source memory across participants, offering further evidence for a role for these regions in episodic memory for meaningful associations. This pattern of results may reflect the myriad of changes in brain structure and function during development (Johnson, 2011), and raises interesting questions regarding how these changes manifest across methods measuring brain activation, connectivity, and representations.

## Conclusion

This study presents important theoretical and methodological advances over prior cognitive neuroscientific studies of memory development, because we 1) examined the interplay of semantic and episodic memory, which has barely been examined, 2) analyzed data from both encoding and retrieval fMRI scans, whereas most prior developmental studies have focused on only encoding or only retrieval scan data, and 3) adopted a multivariate analytic approach that has not yet been employed in children. Our analyses provide converging evidence implicating changes in lateral PFC and its pattern of functional connectivity to age-related improvements in our ability to both process and remember congruent information. During development, these PFC regions may play an increasing role in the service of integrating new experiences with our prior knowledge and encoding them into memory. Beyond development, these results offer insights into the important role of semantic control regions in allowing us to flexibility use our prior knowledge to generate and remember meaningful associations.

## Supporting information

## Acknowledgments

Support for this research was provided by National Institute of Mental Health Research Grant MH091109 (to S.G. and S.A.B.). S.A.B. was supported by a Jacobs Foundation Scholar Award. The authors wish to thank the many research assistants who assisted with data collection, as well as the families who participated in the study.

