## Supplementary material for "The more you know: Investigating why adults get a bigger memory boost from semantic congruency than children"

**Supplementary materials**

### I. Methods: Multi-voxel Pattern Analysis

We performed Multi-Voxel Pattern Analysis (MVPA), a method that detects similarities in the patterns, rather than the magnitude, of activation within a brain region or searchlight sphere. An MVPA searchlight analysis was conducted using The Decoding Toolbox (Hebart, Görgen, & Haynes, 2014) on unsmoothed single trial beta values calculated using the least squares single (LSS) approach (Mumford, Turner, Ashby, & Poldrack, 2012), and subsequently smoothed with an 6mm isotropic full-width at half-maximum Gaussian filter for group analyses. Belong and Don't Belong trials at encoding were used to train a linear support vector machine classifier, which was then tested on the data at retrieval. The searchlight analysis was performed within a 3-voxel searchlight sphere. Area under the receiver-operator-characteristic curve (AUC) maps were outputted for each participant, which provides a measure of classification accuracy that is unbiased by unbalanced conditions (i.e., differences in trial numbers for Belong and Don't Belong conditions).

### II. Results

#### 1. Encoding response times

To better understand the congruency benefit, and the age-related increase therein, we then examined RTs at encoding for item hits to better understand the congruency benefit. A mixed-design ANOVA on encoding RTs (**Supplementary Table 1, top panel**) with factors of age group, congruency, and memory (i.e., Source Correct and Source Incorrect trials) revealed a significant main effect of memory ($F[1,140] = 4.75$, $p = .03$, $\eta_p^2 = .03$), with quicker RTs for subsequent Source Correct than Incorrect trials. No other main effects or interactions reached significance. Thus, RTs did not differ between trials judged congruent or incongruent. Notably, there was no main effect of age group and no three-way interaction with age group; thus, age differences in the congruency proportion, as well as the congruency benefit for source memory, were not due to differences in time on task at encoding.

#### 2. Retrieval response times

In addition to examining RTs at encoding, we tested whether congruent associations were retrieved more quickly, and – if so – whether this effect increased with age. A mixed-design ANOVA on retrieval RTs (**Supplementary Table 1, bottom panel**) with factors of age group, congruency, and memory revealed significant main effects of congruency ($F[1,140] = 28.83$, $p < .001$, $\eta_p^2 = .17$), with quicker RTs for Belong than Don't Belong trials, and of memory ($F[1,140] = 70.97$, $p < .001$, $\eta_p^2 = .34$), with quicker RTs for Source Correct than Incorrect trials. The congruency by memory interaction was not significant ($F[2,140] = 1.51$, $p = .22$, $\eta_p^2 = .01$), indicating that the RT benefit for Source Correct vs. Incorrect trials was comparable between Belong and Don't Belong trials. As expected, there was a significant main effect of age group ($F[2,140] = 7.24$, $p = .001$, $\eta_p^2 = .09$), with slower RTs for younger compared to older children ($p = .04$), with a pattern for older children compared to young adults ($p = .05$). There was also an age group by congruency interaction ($F[2,140] = 5.56$, $p = .005$, $\eta_p^2 = .07$), such that older children ($p = .001$) and young adults ($p < .001$), but not younger children ($p = .35$), were quicker for Belong than Don't Belong trials. There was also an age group by memory interaction ($F[2,140] = 12.16$, $p < .001$, $\eta_p^2 = .15$), as older children ($p < .001$) and young adults ($p < .001$), but not younger children ($p = .06$), were quicker for Source Correct than Incorrect trials. Importantly, the three-way interaction was not significant ($F[2,140] = .52$, $p = .60$, $\eta_p^2 = .01$), indicating that age differences in the congruency benefit for source memory were not due to differences in time on task at retrieval.

**3. MVPA results**

To examine whether activation patterns, across phases, differentiated between the processing of congruency and the lack thereof, we trained an MVPA classifier to distinguish between Belong and Don't Belong trials at encoding, and then tested them on trials at retrieval. This analysis, performed without regard for memory performance, revealed several frontoparietal and temporal regions that exhibited above-chance classification for Belong vs. Don't Belong trials at retrieval in a conjunction of all three age groups (**Supplementary Figure 1**). An F-contrast of age group revealed no suprathreshold regions. This finding suggests that activation patterns associated with evaluating item-context associations are reactivated at retrieval, and that this occurs to the same extent in all age groups.

| | | | Younger Children | Older Children | Young Adults |
|---|---|---|---|---|---|
| Encoding | Belong | Source Correct | 2.40 (.03) | 2.41 (.03) | 2.41 (.04) |
| | | Source Incorrect | 2.43 (.03) | 2.44 (.03) | 2.45 (.04) |
| | Don't Belong | Source Correct | 2.35 (.03) | 2.42 (.03) | 2.42 (.04) |
| | | Source Incorrect | 2.39 (.03) | 2.42 (.03) | 2.41 (.04) |
| Retrieval | Belong | Source Correct | 1.87 (.03) | 1.71 (.04) | 1.47 (.06) |
| | | Source Incorrect | 1.94 (.04) | 1.86 (.04) | 1.81 (.06) |
| | Don't Belong | Source Correct | 1.90 (.03) | 1.78 (.03) | 1.64 (.05) |
| | | Source Incorrect | 1.94 (.04) | 1.93 (.04) | 1.91 (.06) |

**Supplementary Table 1**. Encoding and retrieval response times for each age group. SEM denoted in parentheses.

| Contrast | k | Z | x | y | z | Region |
|---|---|---|---|---|---|---|
| Belong > Don't Belong | 217 | 6.30 | -21 | -79 | -13 | Left Fusiform Gyrus |
| | | 3.85 | -27 | -31 | -19 | Left Parahippocampal Cortex |
| | 76 | 4.36 | -60 | -52 | -4 | Left Middle Temporal Gyrus |
| | | 3.56 | -48 | -49 | -7 | Left Inferior Temporal Gyrus |
| | 66 | 4.11 | 33 | -40 | -13 | Right Fusiform/Posterior Parahippocampal Cortex |
| | 62 | 4.24 | 18 | -70 | 53 | Right Superior Parietal Lobule |
| | 45 | 4.69 | -42 | -67 | 17 | Left Angular Gyrus |
| Age Group x Congruency Interaction | 89 | 4.52 | -48 | 47 | 2 | Left Ventrolateral/Rostrolateral PFC |

**Supplementary Table 2**. Univariate effects for Belong vs. Don't Belong trials. Coordinates are listed in MNI space.

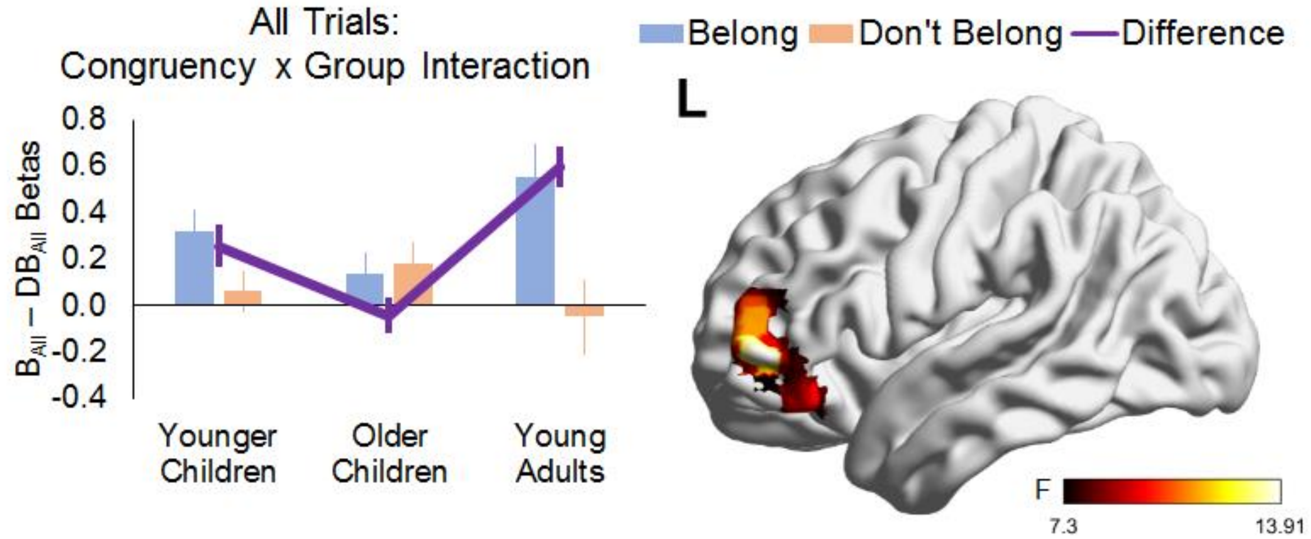


**Supplementary Figure 1**. Left ventrolateral/rostrolateral PFC exhibited a non-linear pattern with age, with greater Belong than Don't Belong activity for all trials in young adults compared to younger children ($p < .001$), and in younger compared to older children ($p < .01$). Post hoc analysis of beta estimates in this ROI indicated that young adults ($p < .001$) and younger children ($p < .01$), but not older children ($p = .45$), exhibited a significant difference between Belong and Don't Belong trials. Moreover, this activation difference was greater in young adults than both younger ($p < .05$) and older ($p < .001$) children; younger children also had a greater activation difference than older children ($p = .01$). Among children, the activation difference between Belong and Don't Belong trials was negatively correlated with age ($r = -0.24$, $p = .01$).

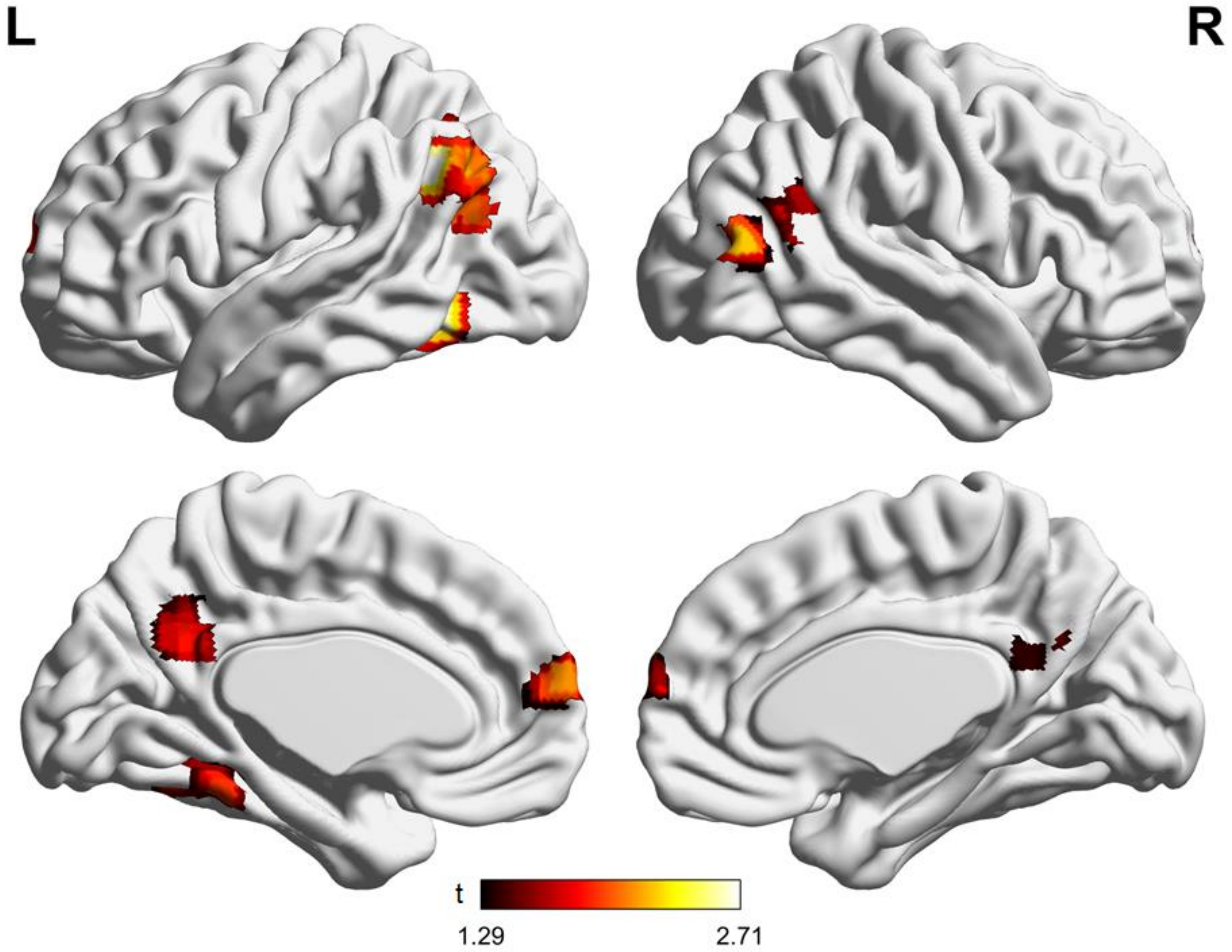


**Supplementary Figure 2**. Regions exhibiting above chance MVPA classification for Belong vs. Don’t Belong trials in a conjunction of the three age groups.
